# Large genetic variability of maize leaf palatability to european corn borer : metabolic insights

**DOI:** 10.1101/2023.04.12.536551

**Authors:** Inoussa Sanane, Stephane D. Nicolas, Cyril Bauland, Frédéric Marion-Poll, Camille Noûs, Judith Legrand, Christine Dillmann

## Abstract

Maize is the most-produced cereal in the world, but its production faces constraints such as parasitic attacks from stemborers. We evaluated the resistance of a core-collection of 18 maize lines by measuring their palatability to European Corn Borer (ECB) larvae fed on maize leaf discs. Using an original consumption test device that takes into account the variability of larvae behaviour, we were able to phenotype the resistance of the 18 maize lines. We matched consumption data to existing enzymatic and metabolomic data that characterized the maize core-collection and identified some metabolites such as caffeoyl-lquinate, trocopherol, digalactosylglycerol and tyrosine that are positively or negatively correlated with the palatability to ECB larvae. Altogether, our results confirm the metabolic complexity involved in the establishment of plant defenses. Metabolic changes associated to leaf palatability mostly concern membrane and cell wall composition. Some of them, pointing-out to the phenylpropanoids pathway, were observed independently of plant developmental pace and plant earliness.

## Introduction

Maize is the leading cereal in terms of production volume, before wheat and rice (24) and plays diverse roles in global agrifood systems, including human alimentation. Worldwide, 6% to 19% of global maize production is lost each year due to insects and other herbivores preying (49). More than 90 insect species are known to feed on cultivated maize (63). Among them European Corn Borer (ECB) *Ostrinia nubilalis* (Hübner), and Mediterranean corn borer (MCB) *Sesamia nonagrioides*, damage maize by boring tunnels within the stems of the plant. Fodder maize plots infested by the European Corn Borer can show up to 80% of plants and 40% of cobs damaged (7), while a single larvae per plant may cause 6% loss in an average grain yield on maize hybrids (9).

In the course of the evolutionary arms race between plants and pests, plants developed many different defense strategies, including physical defenses to minimize the entry of pathogens like cell wall or spines, and biochemical defenses that can be repulsive or toxic (66). They can be constitutive or induced with different resource allocation costs (51). The setting-up of chemical plant defenses is a paradigm of biological complexity. It begins with the exogenous signal perceived from the pathogen and continues with signal perception and signal transduction that may result in the reprogramming of cellular metabolism towards the biosynthesis of secondary metabolites (58). Signal transduction is regulated by hormones (37) and results in a coordinated response mediated by a crosstalk between phytohormones and transcription factors (25, 46).

Indeed, the setting-up of plant chemical defenses has a metabolic cost and mobilizes resources that could have been allocated to other functions like growth or reproduction (31, 32). It may result in trade-offs (28) between different life-history traits. The cost of defenses can affect the carbon-nutrient balance (52), the growth rate (67), or the growth-differentiation balance (61). Those hypotheses are difficult to test (62). Manifestation of detectable trade-offs may depend on the strength of resources limitation or other factors (72). For example, in brown algae, phlorotannins play a role in both primary and secondary metabolism and cannot serve as a reliable indicator (4). However, a recent meta-analysis over a wide range of plants species showed that herbivory reduced growth, photosynthesis and reproduction, but not carbohydrate contents (27).

In maize, resistance to European Corn Borer encompasses both the synthesis of specific antibiosis molecules like DIM-BOA (2,4-dihydroxy-7-methoxy-1,4-benzoxazin-3-one) (13, 59) and changes in the molecular composition of cell wall. In particular, phenolic acids like ferulic or p-coumaric acids may increase leaf toughness (8). More generally, variations in cell-wall phenylpropanoids are associated with resistance to corn borers (29).

One way to measure plant resistance to phytophageous in-sects is to measure leaf disks’ palatability to insect larvae. Such method can also be used to evaluate the antifeedent properties of specific chemicals (1, 40, 47, 48, 56). Feeding preference tests are efficient screen-tests to select resistant plant varieties (18, 64). For example, those methods have been used to measure the palatability of Brassicae plants for *Microtheca punctigera* larvae (44). They allowed for the identification of rice varieties resistant to the lepidoptera *Cnaphalocrocis medinaiset* (6).

Prime to the identification of specific traits associated to resistance or tolerance, it is necessary to evaluate the extent of genetic variability for this trait within the plant species/genetic group using a small number of varieties that represent the genetic diversity within a species or a collection (11, 22). In maize, the evaluation of a panel of 85 inbred lines representing the diversity of the varieties cultivated in Europe allowed for the identification of specific inbreds resistant to *Sesamia nonagrioides* and *Ostrinia nubilalis* after artificial infestation (42), but also for the identification of inbreds able to maintain the plant yield despite pest pressure (43).

In the present study, we used a core-panel of 18 maize inbred lines chosen to represent the genetic diversity of maize varieties cultivated in Europe and North America (10), but also a range of variation for the resistance to European Corn Borer (3, 69) and for cell-wall digestibility (23, 71). Most of the inbred lines from the panel were already shown to present a wide genetic diversity for a large set of physiological, enzymatic and metabolic data (17). We used an original consumption test (56) to measure the genetic variability of leaf-disks palatability to ECB larvae. Making use of the availability of this large dataset, the objectives of the present paper were (i) to assess the amount of genetic variability of maize leaf palatability to European Corn-Borer, and (ii) to seek for correlations between leaf palatability and metabolic or physiological traits that characterized the inbred lines.

## Methods

### Insects rearing

*Ostrinia nubilalis* Hbn.eggs were obtained from Bioline AgroSciences (France). Hatched larvae were maintained in Petri dishes on an artificial diet (1.32l water, 27g agar powder, 224g corn flour, 60g dried yeast, 56g wheat germ, 12g L-ascorbic acid, 4g vitamin mixture and minerals (Réf.0155200), 0.8g chlortetracycline, 2g hydroxybenzoic acid methyl, 1.6g sorbic acid and 4g benzoic acid), under 16 :8 (light: dark) photoperiod at 70% humidity and at 26°C. Second instar larvae (10 days old) were used for the feeding bioassays.

### Plant material: core collection

The plant material comprised 18 maize inbred lines (Table 1). Thirteen of them belong to a core-panel of 19 lines representative of the genetic diversity of modern varieties cultivated in North-America and Europe (10, 14). Those 13 lines were previously characterized at two developmental stages for their variability for central carbon metabolism enzymes activities, metabolites concentrations and a set of physiological traits (17). Among them, two inbred lines (B73, Mo17) are already known for their sensitivity to pyralids attacks (41, 69). The panel was completed with three inbred lines (F66, F271, CM494) exhibiting differences for cell wall digestibility (23, 71), and two inbred lines (F618, F918) known for their tolerance to pyralids attacks (3). All the lines are maintained in the *Centre de Ressources Biologiques INRAE des lignées de maïs* at Saint Martin de Hinx, France. Female flowering time (FFT, TO:0000359 from the Plant Ontology (68)) was predicted by combining data from (10) and yearly measurements at the INRAE field station from Saint Martin de Hinx, France (see Supplementary Methods S1). It was measured in days after sowing. As shown in Table 1, the 18 inbred lines belong to four different maize genetic groups and show a wide range of flowering time variation.

**Table 1.** Description of the maize panel. Pedigree and genetic group of each 18 inbred lines from the panel. Genetic groups are Corn Belt Dent (CBD), European Flint (EF), Northern Flint (NF) or Stiff-Stalk (SS). Lines in **bold** were selected for their tolerance to pyralids attacks. The average female flowering time *FFT* is expressed in days after sowing. The letter above indicates the sowing group for feeding bioassays (L = Late; SL = Semi Late; SE = Semi Early; E = Early).

| Line | Pedigree | Group | FFT |
| --- | --- | --- | --- |
| F64 | Argentina PI 186223 | EF | 79<sup>L</sup> |
| SA24U | Pop-corn | CBD | 77<sup>L</sup> |
| HP301 | Supergold Pop-corn | EF | 76<sup>L</sup> |
| **F918** | **F618 x F630** | SS | 75<sup>L</sup> |
| B73 | Iowa Stiff Stalk synthetic BSSSC5 | SS | 74<sup>L</sup> |
| Lo32 | Isola Basso | EF | 74<sup>L</sup> |
| Mo17 | CI187-2 x C103 | CBD | 73<sup>SL</sup> |
| MBS847 | Pioneer 3901 | CBD | 71<sup>SL</sup> |
| Lo3 | Nostrano dell'Isola | EF | 71<sup>SL</sup> |
| **F618** | **(A166 x B37) x B37** | SS | 70<sup>SL</sup> |
| NYS302 | Black Mexican | NF | 66<sup>SE</sup> |
| C105 | Purple Flour #626 x Ohio Early Yellow Dent inbred #25 | NF | 66<sup>SE</sup> |
| F252 | F186 x Co125 -8.2.3.3.4 | CBD | 63<sup>E</sup> |
| F271 | Co125 x W103 (19.3x8.5) -4.4.1.1.1.1 | CBD | 62<sup>E</sup> |
| Cm484 | (Canada-Morden-1989)200-2-1 |  | 62<sup>E</sup> |
| F2 | Lacaune -2.9.1.1.3 | EF | 61<sup>E</sup> |
| F66 | Sost -15.8 | EF | 61<sup>E</sup> |
| ND36 | Manitoba Yellow Flint | NF | 59<sup>E</sup> |

### Plant material: growing conditions

To compare the inbred lines at the same developmental stage, flowering time was used to constitute four different sowing groups (Table 1) and to plan four different sowing dates by group, in order to constitute at least three blocks with all maize lines sampled at the same developmental stage. Altogether, sowing were realized between october 1st 2019 and november 1st 2019 (Supplementary Methods S1). For each sowing date, six seeds per line were pre-germinated on sowing plates until 3-4 visible leaves. Then each plant was repoted in 4L pots containing Jiffy® premium substrate and deposited on a shelf that contained plants from the 18 inbred lines at the same developmental stage and constitute a replicate for the feeding bioassays (Fig 1a). Plants were cultivated in a greenhouse under 16:8 (light: dark) photoperiod with 70% humidity and a temperature comprised between 21 and 24°C. Pots were watered two times a week. The position of the pots in the shelf was randomized.

**Fig. 1.**
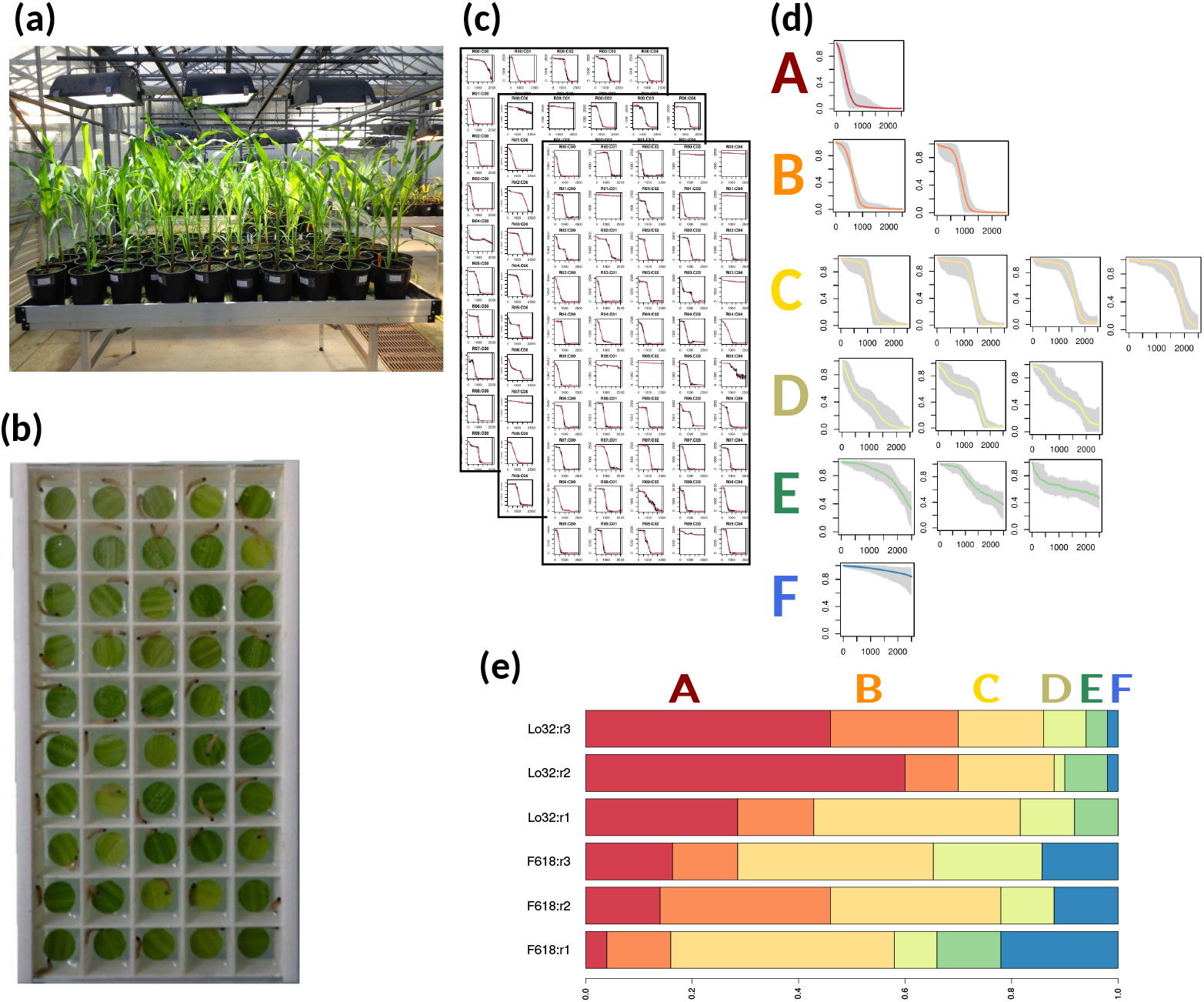
Overview of the larvae feeding bioassays. **(a)** Plants from the 18 inbred lines of the panel were grown in a greenhouse. Delayed sewing allowed to sample plants at the same developmental stage for each replicate. **(b)** For each inbred line/replicate, 50 leaf discs were arranged into a 50-cells plate. A single L2 larva was deposited into each cell at the beginning of the monitoring. **(c)** The consumption of 50 leaf discs by L2 larvae was monitored during approximately 24h. Image analysis allowed to measure the proportion of leaf discs consumed at each time step. **(d)** Clustering methods allowed to classify the consumption curves into six insect larvae behavioural groups, from consumers (group A) to non-consumers (group F). **(e)** The output was the proportion of each behavioural type in each replicate. As an example, the three replicates from the two most extreme lines *F* 2 and *Lo*32 are shown

### Feeding Bioassays

For enabling data comparison with metabolic and physiological data collected by (17), we chose to sample the vegetative developmental stage between GRO:0007011 (tassel initiation) and GRO:007013 (ear initiation) from the Cereal Plant Development Ontology (68). This corresponds to plants having between 5 (V5) and 7 (V7) visible leaf collars.

For each inbred line, 4 × 50 leaf disks were tested. 1 cm diameter leaf disks were punched from the 6th leaf of 3 plants of approximately the same developmental stage. Each leaf disk was quickly placed upon a 5 mm layer of 1% agarose within a cell from a 5 × 10 cells grid. Subsequently, one L2 instar larva was placed into each cell and its feeding activities were monitored during 48h. As our experimental setup allowed us to test simultaneously only 6 grids, the experiment was run as 4 repetitions x 3 batches x 6 inbred lines x 50 leaf disks (of 1 inbred line) (see Supplementary Methods S1).

Leaf disc consumption by L2 larvae was monitored for each individual cell for 48h and using the recording system described by (56). Image stacks were analyzed using the plugins *RoitoRoiArray* and *Areatrack* developed in the laboratory (56) to run under the software Icy (19). These plugins were used respectively to delimit the position of each cell on the images and to evaluate the surface of each leaf disk across time. The measures (pixel per minute) for each leaf disc in each well were exported in an Excell spreadsheet. Data are converted into CSV flat files and further analyzed with the R software (50).

Among the four blocks, one was discarded because larvae were not at the right L2 stage when the plants where at the correct V5-V7 developmental stage.

### Statistical analyses of feeding bioassays: behavioural types

As in (56), data analyses were conceived as a two-stage procedure. The first stage consisted in describing the variability of individual larvae feeding behaviours and classifying them into six behavioural types. At the end of this stage, each replicate of each inbred line is characterized by a distribution of behavioural types among the 50-wells. The full procedure and corresponding R scripts are available online (54).

Briefly, the R scripts generates pdf files representing consumption curves for each larva in each well (Fig. 1c). Different larvae are not expected to have exactly the same behaviour even when submitted to the same conditions. A non-supervised hierarchical classification of the 2700 individual consumption curves corresponding to the 3 blocks of the 18 inbred lines was realized using the SOTA algorithm (33). It ended-up with 14 clusters identified by a letter from *a* to *n*. To reduce the number of groups, each cluster was characterized by summary statistics like the average times T20, T50 and T80 to consume 20, 50 or 80% respectively of the leaf disc area, or the total consumption. Based on those summary statistics, the Kmeans algorithm (30) along with some manual grouping were used to group the 14 clusters into six ordered feeding behavioural types, named from *A* to *F*. Average feeding behavioural profiles and their range of variation are presented on Fig. 1d.

- *A, B, C* behavioural types mainly differ in the existence or not of a lag-time before consuming, and in the length of the lag-time. Generally, the leaf is fully consumed at the end of the experiment. *A* types correspond to fast consumers.
- *D, E, F* behavioural types are reluctant to consume. Consumption rate is low. Generally, the leaf is not fully consumed. Type *F* larvae are non-consumers.

At the end of the process, each consumption curve, corresponding to a single well in a single plate is attributed a behavioural type, from *A* to *F*. For each inbred line *i* and each block *j*, the distribution of behavioural types can be counted. We called 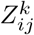 be the number of observations of behavioural type *k* ∈ {*A, B, C, D, E, F*} from inbred line *i* in replicate *j*. Fig. 1e shows three examples of the *Z*_*ij*_ distribution, corresponding to the three replicates from inbred line Lo32 and the three replicates from inbred line F618.

### Statistical analyses of feeding bioassays: AFratio

Rigorously, we could use multinomial regression to test whether line or block change the behavioural distribution. However, the experimental set-up, with three replicates per inbred line lacks of power. Instead, we proposed to build quantitative statistics to measure consumption behaviour by reducing the behaviours into two classes: *consumers* versus *reluctants*:

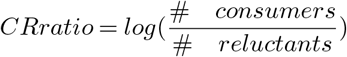

There are many possible combinations to group six classes (A to F) into two (consumer, reluctant). Among all possible combinations, we decided to choose the one that allowed for the best discrimination between the inbred lines. For each possible combination, we ran a linear model

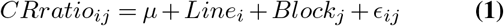

and recorded the adjusted *R*2 for the model, as well as the heritability of the line effect

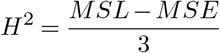

where *MSL* is the mean square associated with the Line effect, *MSE* is the error mean square and 3 is the number of blocks.

Results are detailed in Supplementary Methods S2. The most discriminant combination was the AFratio, *i*.*e* the log-ratio of the proportion of *A*-type fast consumers over the proportion of *F* -type non-consumers:

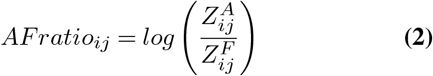

The anova model (eq 1) was used to compute the mean AFratio for each inbred line, 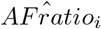, as well as confidence intervals. Comparison of inbred lines means were performed using Tukey Honest Significant Difference tests (70). Pearson’s correlation coefficient between AFratio and Flowering time (FFT) was also computed.

### Correlations with plant metabolism and physiology

Thirteen inbred lines from our panel were characterized at two developmental stages for a set of physiological, enzymatic or metabolic data. We used the data available as Supplementary Dataset1 from (17) to compute the average value for each inbred line. In (17), the vegetative stage *V* was considered as plants with 7 to 8 visible collar leaves. It roughly corresponded to the ear initiation stage (GRO:0007013 from the Cereal Plant Development Ontology) and is comparable to the developmental stage used in the present study. As in the present study, samples from the sixth leaf were used for enzymatic, metabolomic and physiological analyses. The second developmental stage was 15 calendar days after silking (15DAS). It corresponded to the blister stage (GRO:0007030). This stage is initiated when significant starch accumulation begins, approximately 12-17 days after pollination.

Altogether, the dataset comprised enzymatic activity (Vmax) from 29 enzymes from central carbon metabolism and relative concentration (*nmol*.*mg*^−1^*leaf FW*) from 155 metabolites at two developmental stages (V and 15DAS). It also comprised the measurement of Yield, kernel number and Thousand Kernel Weight at maturity, as well as dry matter content, C/N ratio, and the C, N and nitrates content at the two developmental stages (V and 15DAS).

Among the 383 traits measured, 228 were variable between the 13 lines of our panel. 139 traits showed a quantitative variation within our panel. Eighty-nine traits showed a qualitative variation (presence/absence or no more than three different abundance values). Among those 89 traits, 27 were present or absent in a single inbred line and were subsequently discarded. The 62 remaining traits with presence/absence were treated as qualitative variables. For each trait *l*, the abundance was transformed into a qualitative variable 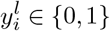. Its relationship with AFratio was analyzed with a linear model :

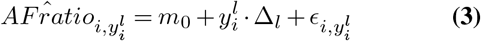

where *m*_0_ is the mean *AFratio* among the inbred lines where the trait is absent, and Δ_*l*_ is the average effect of the presence of trait *l*.

For the 139 traits with quantitative variation within the panel, Pearson’s correlation coefficient with AFratio and FFT was computed, as well as the corresponding pvalue. Traits with a *pvalue <* 0.05 were retained as associated. A Principal Component Analysis was run to explore the correlations between quantitative traits that were found significantly associated with AFratio.

## Results

We used an original high-throughput design for feeding bioassays (56) to measure genetic variation of maize leaf palatability to second instar larvae from the european cornborer *Ostrinia nubilalis* Hbn within a maize inbred lines core-panel. The maize panel covered the main maize genetic groups that represent the diversity of maize varieties cultivated in Northern America and Europe (10). As shown in Table 1, the panel presented a wide range of variation for flowering time between ND36, that flowers 59 days after sowing, and F64 that flowers 79 days after sowing.

### Classifying feeding behaviours

Maize leaf palatability was assessed during the vegetative growth stage, when plants exhibit between 5 and 7 visible leaf collars. Delayed sowing dates were used to sample plants from the different inbred lines at the same developmental stage. Feeding bioassays consisted in measuring the consumption of leaf discs from the sixth leaf by second instar pyralids larvae. Instructions for building-up the feeding consumption bioassays device are freely available (55). Image analysis was performed using plugins embedded into the image software Icy (19). R scripts for statistical analyses were deposited in (54). Fig. 1 gives a general overview of the process. Clustering methods were used to classify individual consumption curves into six ordered behavioural types, named from *A* to *F* that captured both differences between leaf samples and behavioural differences between larvae. Fig. 1d, shows the percentage of intact leaf disc as a function of time for each behavioural type. Clearly *A* types are *consumers* that feed fast and consume all the leaf disc, while *F* types are *reluctants* that hardly feed on the leaf disc. In between, *B* to *E* behavioural types are intermediate. *B* and *C* mainly differ from *A* by the existence of a lag-time: larvae wait before feeding. *D* and *E* mainly differ from *F* by the fact that at least part of the leaf disc is consumed at the end of the experiment. They differ from *A, B* or *C* by the consumption rate, which is always lower. The same range of variation of behavioural types were observed in (56), where larvae were confronted to leaf discs from a single maize variety treated with different concentrations of antifeedant molecular compounds. Here, in addition to the variability of larvae, **the variability of behavioural profiles reflects natural variations for palatability between sampled leaf discs**. Those differences may come either from growing conditions or from differences between the inbred lines.

### Assessing genetic differences for maize leaf palatability

In order to assess genetic differences between lines, the distribution of behavioural types was established for each inbred line and each replicate by counting-out the number of leaf discs exhibiting the different behavioural types. Fig. 1 shows the results for the three replicates from inbred lines Lo32 and F618. Clearly, there were variations between replicates. However, the proportion of *A* behavioural type is always high in Lo32 and low in F618, while the proportion of *F* behavioural type is always low in Lo32 and high in F618. Fig. 2a shows the average distribution of behavioural types for each inbred line of the core-panel. The average proportion of *A*-type behaviours ranges from 12% in F618 to more than 40% in Lo32 and exhibits a quantitative variation within the panel. The average proportion of *F* -type behaviours decreases with the average proportion of *A*-type behaviours. It ranges from 18% in F618 to 2% in Lo32. Hence, genetic differences between lines at least partly drive the observed differences between leaf discs.

**Fig. 2.**
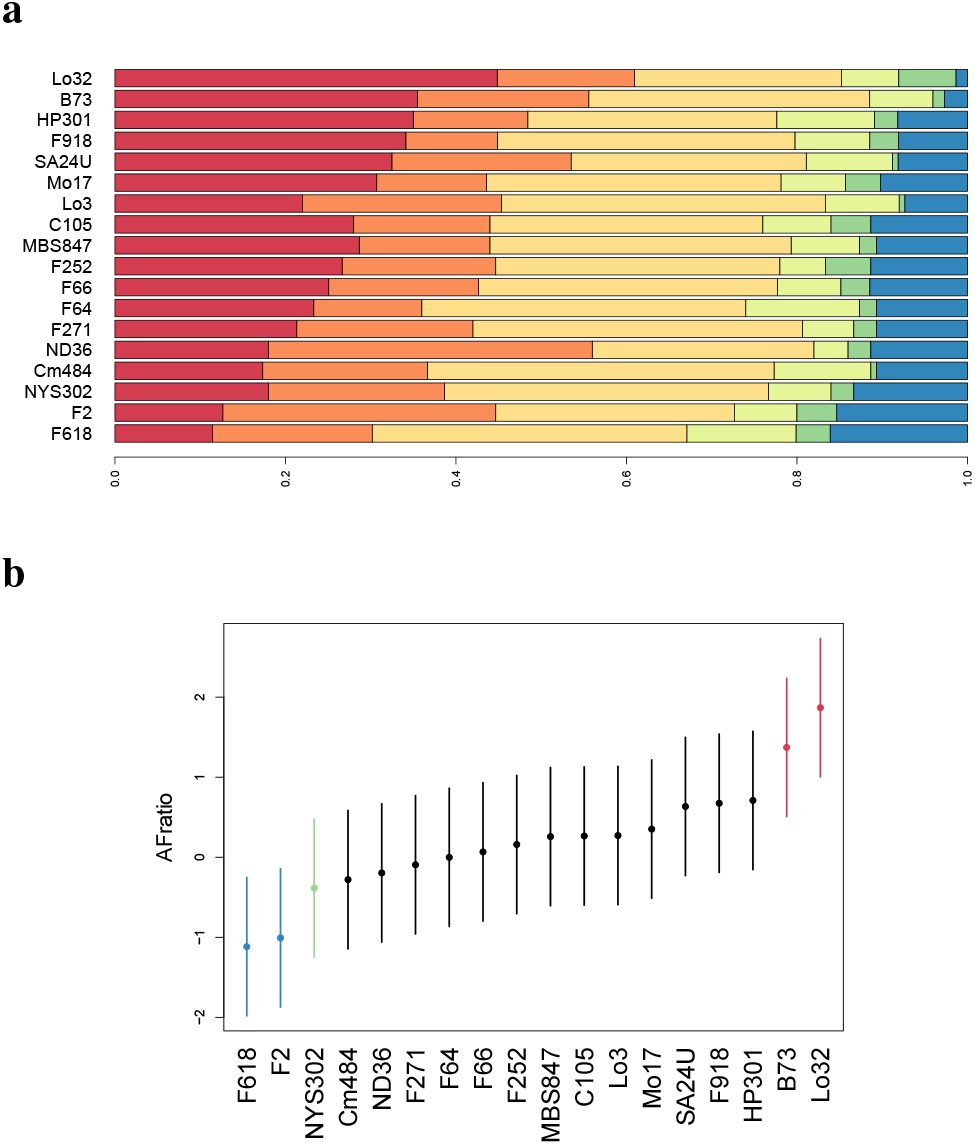
Feeding bioassays. **a.** Average distribution of behavioural types for each of the 18 inbred lines from the core-panel. Colors are the same as in Figure 1. Red is the proportion of A types, and blue the proportion of F types. **b** Range of variation for the AFratio. Dots indicate the mean. Lines the 95% confidence interval around the mean. Maize lines are ordered according to their mean AFratio. Colours highlight groups of lines with significant differences.

To test for differences between inbred lines, AFratio was computed as the log-ratio of the number of leaf discs attributed to the *A* behavioural type to the number of leaf discs attributed to the *F* behavioural type (eq 2). The AFratio was shown to be the most discriminant log-ratio among all possible ratios (Supplemental Methods S2). An analysis of variance taking into account the inbred line and the block effects (eq 1) showed that both effects were significant: the line effect pvalue was 0.0015 and the block effect pvalue was 0.007. Indeed, blocks 2 and 3 tended to have a higher AFratio than block 1 (Fig. 7 from Supplementary Methods S2). The between-line heritability was *H*2 = 0.40. Fig. 2b shows the mean AFratio and its confidence interval estimated for each inbred line of the panel. AFratio ranged from −1 (F-types were two times more frequent than A-types) in F618 and F2 to +2 (A-types were seven times more frequent than F-types) in Lo32. In B73, A-types were on average two times more frequent thant F-types. Inbred line NYS302 was close to F618 and F2 and significantly different from B73. Despite the lack of power to detect significant differences, AFratio showed a continuous variation between lines that reflects genetics differences of maize leaf palatability to European cornborer.

Fig. 3 shows the positive correlation between AFratio and flowering time (r=0.55, pvalue=0.017). Inbred lines that flower earlier seem to be less attractive to pyralids, with an excess of *F* behavioural types. However, notice that the four extreme lines for AFratio : F2, F618, B73 and Lo32 strongly depart from the regression line, suggesting that flowering time is not the sole determinant for the variation of AFratio.

**Fig. 3.**
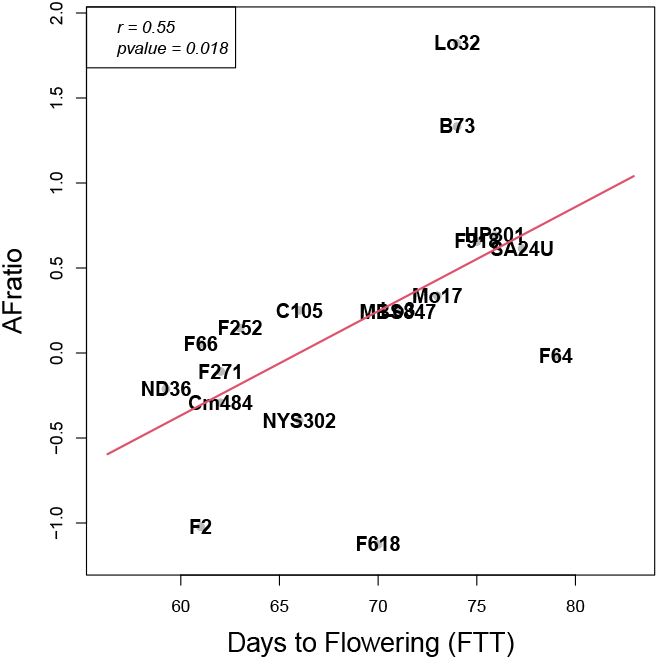
Correlation between Flowering time and AFratio. The scatterplot shows the relation between the average AFratio estimated for each inbred line and its average female flowering time FFT. The red lines corresponds to the regression line.

### Correlation with metabolic and physiological traits

Thirteen inbreds lines of the panel were thoroughly characterized for a large set of enzymatic, metabolic and physiological traits (17) at two developmental stages : vegetative (V) and grain-filling (15DAS). The vegetative stage, around rapid stem elongation and ear initiation was the same as the one targeted in the present study for feeding bio-assays. We took the opportunity of the availability of the data to explore the relationships between maize leaf palatability measured by *AFratio* and metabolic or physiological characteristics of the inbred lines.

Among the 62 qualitative traits that were either present or absent and where analyzed by anova using (eq 3), four metabolites and one amino-acid concentrations showed a significant effect on AFratio and were reported in Table 2. They all belong to the phenylpropanoid pathway (3). Notice that the average effect associated with the presence of the compound was important and corresponded to almost half of the differences in leaf palatability between the two most extreme lines. Presence/absence of the compound defines two groups of lines, one always comprising the less palatable line F2, and one always comprising the most palatable line B73. The presence of Tyrosine, Coumaroylquinate, and To-copherol in the F2 group is associated with a decrease of leaf palatability. The presence of Caffeoylquinate.trans and Caf-feoylquinate.cis in the B73 group is associated to an increase of leaf palatability. Notice that the number of maize lines composing the two groups changes depending on the compound. While the organic acid *Coumaroylquinate* was only present in the three less attractive inbred lines of the sub-panel (F2, ND36, NYS302), the effect of the presence of the other compounds seems to be less clearcut, suggesting that the modulation of leaf palatability has complex mechanisms.

**Table 2.** Association between AFratio and qualitative traits. Δ is the average effect of the presence of the trait on AFratio (eq 2). Pvalues are between brackets. When inbred line names are *emphasized*, this means that the compound is absent. Otherwise, list of the inbred lines where the compound is present. Abbreviated traits names are the same as in (17).

| Trait | $\Delta$ (pvalue) | Lines |
| --- | --- | --- |
| DAS.Tyrosine | -0.96 (0.011) | F2, F64, MBS847, ND36, NYS302 |
| V.4.Caffeoylquininate.trans | 0.85(0.045) | F2, *F252*, *F64* |
| V.3.Caffeoylquininate.cis | 0.79(0.050) | B73, C105, Lo32, MBS847, MO17 |
| DAS.Coumaroylquininate... | -1.11 (0.011) | F2, ND36, NYS302 |
| DAS.Tocopherol | -1.61 (0.031) | Lo32, SA24 |

Among the 139 traits with a quantitative variation within the sub-panel, 17 were significantly correlated to AFratio, while 13 of them were also significantly correlated to flowering time (Table 3). Fig. 4 shows the correlations between those 17 traits and flowering time in the form of a principal components’ analysis. The first PCA axis explains 62% of the total inertia and separates inbred lines according to their flowering time. It is mainly driven by differences between the late Lo32 and SA24U and the early F2, that were also the most extreme for leaf palatability. Traits correlated to PCA axis1 were mainly traits measured at a late developmental stage (15DAS) during grain filling. Early inbred lines are associated with higher activity of enzymes involved in carbon fixation and nitrogen assimilation 15 days after silking (15DAS). This results in a higher percentage of nitrogen and nitrates, and a lower C/N ratio. For those traits, causal relationships with AFratio are difficult to disentangle from a pleiotropic effect of phenology. The second PCA axis explains 8% of the total inertia and separates lines that belong to the group with a high AFratio (C105, MO17, B73) from lines that belong to the group with a lower AFratio (SA24U, HP301, ND36). Traits correlated to PCA axis 2 were mainly traits measured during the vegetative development (V). In particular, PCA axis 2 shows a strong positive correlation with *caffeoylquinate*, an organic acid involved in the biosynthesis of lignin (phenylpropanoids) that belong to plant secondary metabolism.

**Table 3.**
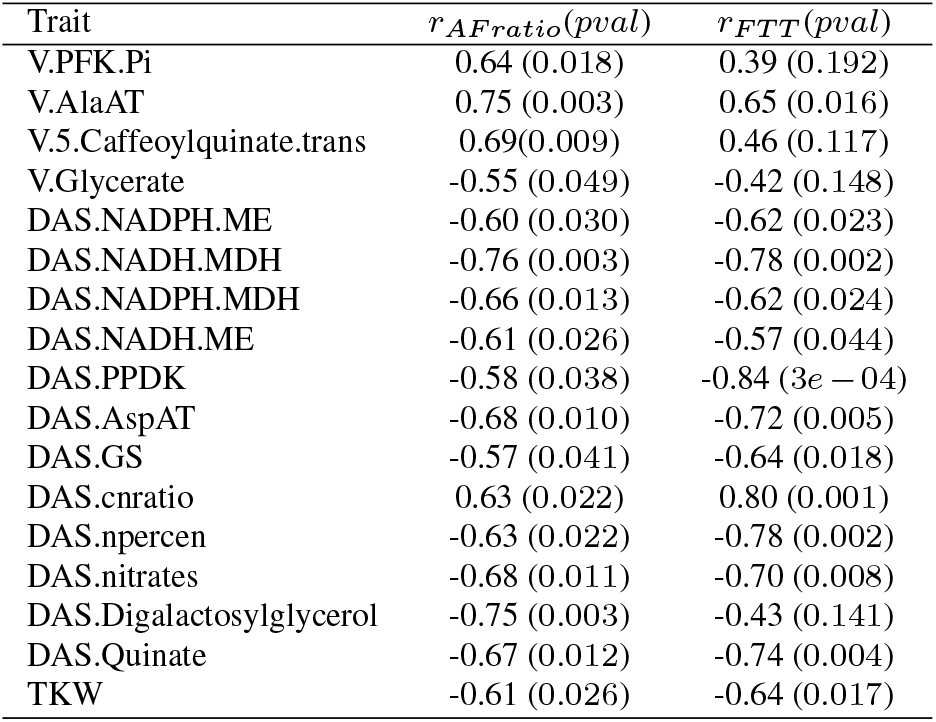
Correlation between AFratio, FTT and quantitative traits. For each trait, Pearson pairwise correlation coefficient with AFratio (eq 2) and flowering time (FTT), respectively. Corresponding pvalues are given between brackets. Traits names were the same as in (17).

| Trait | $r_{AFratio}(pval)$ | $r_{FTT}(pval)$ |
| --- | --- | --- |
| V.PFK.Pi | 0.64 (0.018) | 0.39 (0.192) |
| V.AlaAT | 0.75 (0.003) | 0.65 (0.016) |
| V.5.Caffeoylquininate.trans | 0.69(0.009) | 0.46 (0.117) |
| V.Glycerate | -0.55 (0.049) | -0.42 (0.148) |
| DAS.NADPH.ME | -0.60 (0.030) | -0.62 (0.023) |
| DAS.NADH.MDH | -0.76 (0.003) | -0.78 (0.002) |
| DAS.NADPH.MDH | -0.66 (0.013) | -0.62 (0.024) |
| DAS.NADH.ME | -0.61 (0.026) | -0.57 (0.044) |
| DAS.PPDk | -0.58 (0.038) | -0.84 (3e - 04) |
| DAS.AspAT | -0.68 (0.010) | -0.72 (0.005) |
| DAS.GS | -0.57 (0.041) | -0.64 (0.018) |
| DAS.cnratio | 0.63 (0.022) | 0.80 (0.001) |
| DAS.npercent | -0.63 (0.022) | -0.78 (0.002) |
| DAS.nitrates | -0.68 (0.011) | -0.70 (0.008) |
| DAS.Digalactosylglycerol | -0.75 (0.003) | -0.43 (0.141) |
| DAS.Quinate | -0.67 (0.012) | -0.74 (0.004) |
| TKW | -0.61 (0.026) | -0.64 (0.017) |

**Fig. 4.**
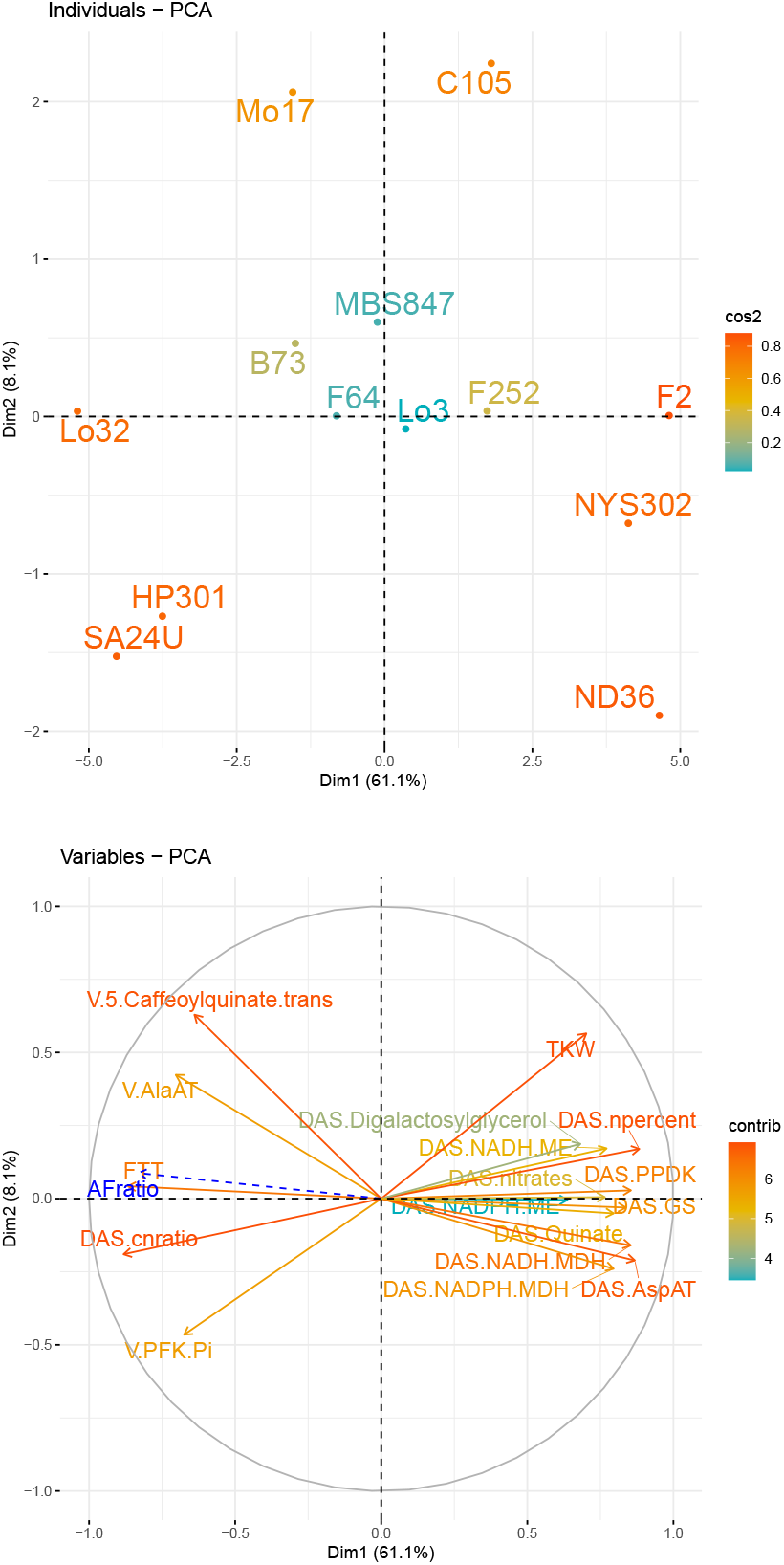
Correlations between associated quantitative traits. Results from the Principal Component Analysis with AFratio as supplementary variable. **Top** Position of the inbred lines in the 2-dimensional space engendered by the first two PCA axes. **Bottom** Correlation circle. The colour code corresponds to the contribution of individuals or variables to the PCA axes.

Altogether, among the 201 variable traits, 22 of them were found associated to AFratio at the 5% level, which is two times more than expected under the null hypothesis because of multiple testing. Indeed, the False Discovery Rate computed from the observed distribution of the pvalues is around 0.45. The functional annotation of all metabolites from Table 2 and Table 3 was achieved using the MetaCyc (16) and KEGG (36) databases and presented in Supplementary Data S3. It shows that the associated traits were enriched in traits linked with the phenylpropanoid pathway.

## Discussion

We used a new feeding consumption test (56) to evaluate maize leaf discs’ palatability to European Corn Borer larvae within a core panel of 18 temperate maize inbred lines. The objectives were to assess the extent of genetic variability for palatability within the panel, and to link those variations to plant metabolism.

Most consumption tests characterize leaf consumption through time by a single instant parameter like the time to consume half of the leaf disk (20, 44, 57). Those tests typically lack of power for two reasons. First, they fail to take into account the natural variability of individual larvae behaviours (6). Second, because larvae may change their behaviour through time (34) and (Fig1d). Our experimental set-up bypasses both drawbacks. First, it allows for the observation of the feeding behaviour of a large number of individual larvae (50) within each biological replicate. Second, instead of summarizing the behaviour by an instant or average value, it proposes an original method to classify individual consumption curves into ordered feeding behavioural types. In this study, we observed six main feeding behaviours that go from the immediate consumption of the whole leaf disc (A type) to the absence of consumption (F type). Intermediate behaviours correspond to lag-time before consumption (B and C types) or slower consumption rates (E type) with breaks (D type). In a previous study, we demonstrated the link between larvae behavioural type and leaf palatability by using antifeedant molecules (56). Here, we observe the same kind of variability for larvae behaviour when confronted to different maize inbred lines. The distribution of the different feeding behaviour types within a biological replicate takes into account natural variability between individual larvae and measures the average palatability of the sample. We propose here a quantitative measure of leaf palatability, named AFratio, and computed as the log-ratio of the two most extreme behaviours, A and F. **We found genetic variability for AFratio between inbred lines within the panel with a broad-sense heritability around** *H*2 = 0.40 **and confirmed the interest of the method for consumption tests**.

While our data clearly show genetic variation for leaf palatability to ECB, we only have indirect evidence concerning the link between leaf palatability and the setting-up of plant defenses in the field. However, our results can be compared to experimental evidences concerning tolerance/sensitivity to ECB. Classically, tolerance to ECB is assessed through the measurement of plant damages after artificial field infestation. Our panel comprised a few inbred lines for which tolerance/sensitivity to ECB have already been assessed in field experiments. The tolerant line *F* 618 (3) is the less palatable from our panel while the sensitive line *B*73 (41, 69) is amongst the lines with the highest AFratio (Fig 2). *Mo*17 is reputed to be sensitive and stands in the top 6 inbred lines with the highest AFratio. In the same line, the relative ordering of the lines *B*73 (36% of A types and 3% of F types), *HP* 301 (35% of A types and 8% of F types) and *Mo*17 (31% of A types and 11% of F types) is similar to the one obtained by measuring *S. frugiperda* Smith larvae growth rates on leaf disks (35). However, the link between leaf palatability and the amount of plant damages in the field stays complex. The inbred line *F* 918 shows a moderately high palatability while it was derived from *F* 618 and selected for tolerance. **Altogether, our new feeding consumption test allowed us to classify the panel inbred lines for leaf palatability. Lines** *Lo*32 **and** *B*73 **were the most palatable, and lines** *F* 618, *F* 2 **and** *NY S*302 **the less palatables**.

Interestingly, the link between leaf palatability and earliness is not straightforward. There is a moderate positive linear correlation between earliness and AFratio (*r* = 0.55, 1 *pvalue* = 0.018, Fig 3). Fast development tends to be associated with a higher level of defenses when measured during the vegetative plant stage, in contradiction with the *growth or defend* trade-off (31). However, Late lines exhibit a wider range of variation for leaf palatability and comprise both the tolerant poorly palatable *F* 618 and the sensitive highly palatables *B*73 and *Lo*32. Such patterns can be explained by gain or losses of metabolic functions due to random genetic drift or selection history (remember that *F* 618 have been selected for tolerance to ECB (3)). The relatively high palatability of early maize inbreds could be explained by local adaptation between plant and insect phenology: in environments favorable to the culture of early maize varieties, insect phenology leads to earlier attacks and resulted in the selection of plant lines able to mobilize their defenses at earlier developmental stages. This hypothesis could have been tested by setting-up leaf consumption tests at different plant developmental stages.

In maize, there is a long standing literature about genetic variability for plant defenses against herbivores that concerns both induced and constitutive defenses. For example, maize inbred lines differ for the volatile compounds emissions induced by injection of *Sprodoptera littoralis* regurgitant (21). Genes involved in the phenylpropanoid pathway were shown to be polymorphic (2). Within the phenypropanoid pathway, QTLs were found for stem-wall hydroxycinnate contents like p-coumaric or ferulic acids (38), but also for resistance to lepodiptera *Spodoptera frugiperda* Smith and coleoptera *Sitophilus zeamais* (5). Using a MAGIC population of 408 recombinant inbred lines, (39) showed that a greater concentration of p-coumaric acid was associated to a higher resistance to corn-borers, measured by tunnel length in infested plants, and also a lower yield. Altogether, those studies evidence the metabolic complexity of plant defenses and pinn-point the central role of phenylpropanoid pathway (58).

Here, we benefited from the availability of the metabolomic and enzymatic characterization of 13 of our 18 core panel inbred lines (17) to investigate the metabolic bases of maize leaf palatability. Among the 201 variable metabolic traits, only 22 were found significantly associated to variations in maize leaf palatability. Interestingly, those 22 metabolic traits were clustered into a small number of metabolic pathway according to https://metacyc.org: chlorogenic acids pathways, chorismate-tyrosine pathway, malate metabolism, hydroxilated fatty-acids pathway, all involved in the establishment of plant defenses. Besides, maize leaf palatability is associated with a high C:N ratio and a low concentration of nitrogen and nitrates, as well as with a low yield. In tomato, C:N ratio was considered as a good indicator of secondary compounds concentrations, especially those involved in the chemical defenses (52).

Digalactosil-glycerol is a glycolipid specific from plant plasma membrane possibly associated to host-pathogens interactions (65). A high level of this compound or its pre-cursor glycerate (Fig S8) is associated with a low palatability, while a high enzymatic activity of phosphofructokinase (PFK), which mediates carbon allocation to pentose-phosphates, sucrose or hydroxilated fatty acids is associated with higher palatability. Malate metabolism is at a crossroads between gluconeogenesis and the biosynthesis or aromatics amino-acids (15). We found five enzymes from C4 dicarboxilic acid cycle and nitrogen assimilation that lead to tyrosine biosynthesis and were associated to maize leaf palatability (Fig5). A high level of activity for those enzymes is associated with a lower leaf palatability and a higher tyrosine concentration, except for alanine-aminotransferase. Note that alanine-aminotrasferase is at a crossroads between tyrosine and alanine biosynthesis. Chorismate-tyrosine pathway is another regulatory hub that was shown to control vitamine E content in tomato (12). In our study, both tyrosine and trocopherol concentrations were found negatively correlated with maize leaf palatability. Trocopherol is an amphiphilic lipid with vitamine E activity. It protects membranes against oxidative stress with a special role associated with the protection of plant photo-system II (45). Trocopherol biosynthetic pathway modulates salicylic acid accumulation and affects basal resistance against *Pseudomonas siryngae* in the model plant *Arabidopsis thaliana* (60). Finally, we found three metabolites from the chlorogenic acids pathway which concentration was associated to leaf palatability : coumaroyl-quinate, caffeoyl-quinate and quinate. Those metabolites are substrate and products of two successive enzymatic reactions (Fig 5) and possibly linked to trocopherol biosynthesis through quinate degradation. The enzyme that transforms coumaroyl-quinate into caffeoyl-quinate have been identified as p-coumarate 3-hydroxylase (C3H) found in the *ref8* mutant in *Arabidopsis thaliana* (26). *ref8* plants deposit an unusual lignin enriched in p-hydroxyphenyl sub-units and are prone to fungal attacks. It was suggested that phenylpropanoid pathway products downstream of REF8 may be required for normal plant development and disease resistance. **Altogether, our results confirm the biological complexity of the metabolic response associated to plant defenses (58). However, all metabolic changes related to leaf palatability seem to be related to changes in membrane and cell-wall composition**.

**Fig. 5.**
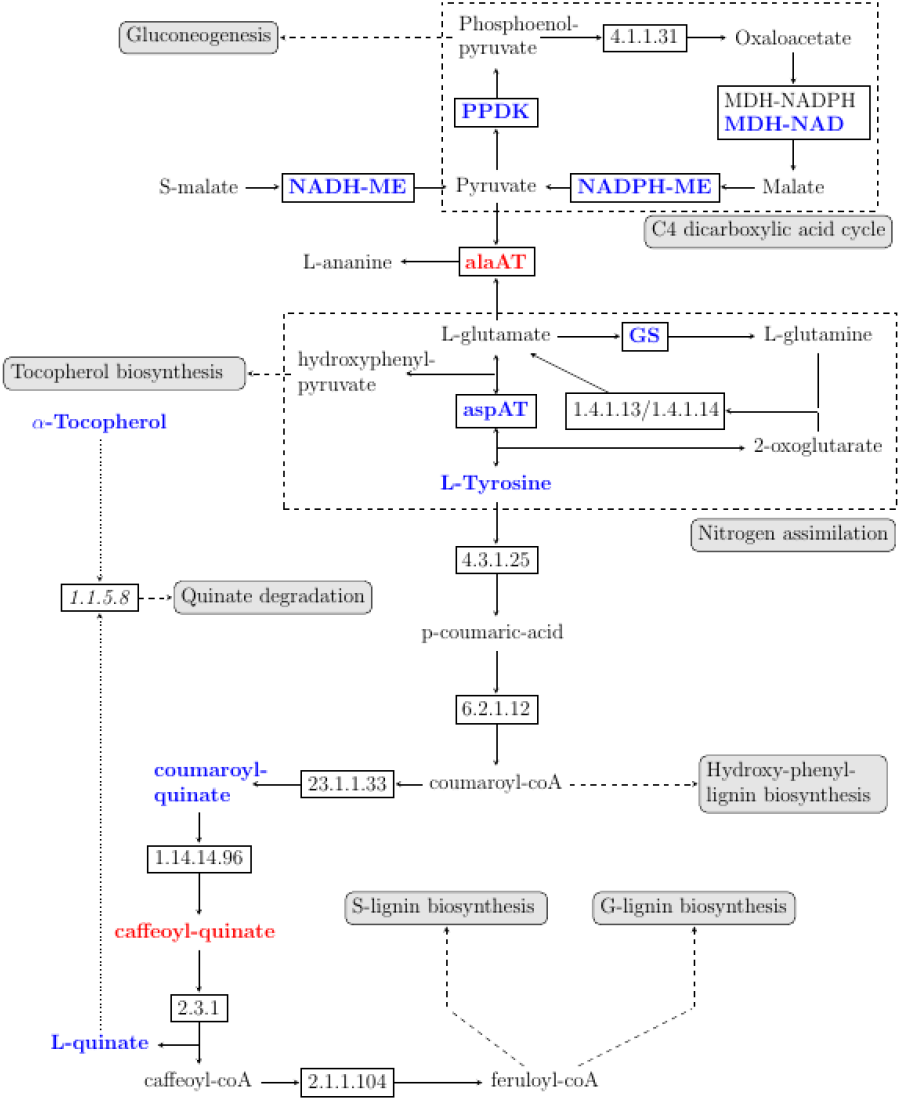
Metabolic pathways associated to maize leaf palatability: phenyl-propanoids. Enzymes (rectangles) are given their EC number or their abbreviated name. Straight lines indicate a direct relation between enzymes and sub-strates/products and arrows the main direction of the reaction. Dashed lines indicate the link to a pathway. Point lines indicate an hypothetical direct reaction. Colors indicate a significant positive (red) or negative (blue) association with maize leaf palatability.

## Conclusions

The original consumption test used in this study allowed-us to highlight genetic variability of leaf palatability to European Corn Borer within a core-panel of maize inbred lines representative of the varieties cultivated in temperate areas. Our results are in accordance with existing data about tolerance/sensitivity of the inbred lines observed in the field. Correlation analyses between leaf palatability and the concentration of metabolites and enzymes points out candidate maize metabolic pathway that could be explored through functional analyses.

## ACKNOWLEDGEMENTS

This study has benefited of a grant from Institut Diversité, Ecologie, Evolution du Vivant (IDEEV) and it was supported by a scholarship from the Islamic Bank of Development to Inoussa Sanane (N° BID: 600033174). We would like to thank INRAE Maize Germplasm Bank at Saint Martin de Hinx CRB INRAE des lignées de maïs, specifically its director Carine Palaffre, for providing us with the INRAE inbred line accessions, and the North Central Regional Plant Introduction Station (NCRPIS) for the non-INRAE inbred line accessions.

## Data availability

The maize lines used in this paper are available upon request from INRAE Maize Germplasm Bank at Saint Martin de Hinx. Data and Rscripts are fully available from the French national platform data.gouv.fr (53).

## Supplementary Data S1: Feeding bioassay experimental set-up

### Flowering time and precocity groups

For each inbred line, data about flowering time came either from (10) or from the yearly recordings of the INRAE field station at St Martin de Hinx, with 10 out of the 18 lines having records for both. In (10), female flowering time (FFLW) was recorded in sum of temperatures while data from St Martin de Hinx (FFT) were recorded in days after sewing. A linear regression was performed on the set of common lines to predict the FFT from the lines that were not measured at St Martin de Hinx. Hence, the flowering time (FFT) presented in Table 1 results either from observation at the INRAE field station or from predictions. FFT data were used to group the maize lines of the panel into four sewing groups, labelled from A to D. All available information is summarized in Table S4

**Table 4.** Flowering time informations.

| Line | FFLW | FFT<sup>a</sup> | LFNB | FFT<sup>b</sup> | sewing group |
| --- | --- | --- | --- | --- | --- |
| ND36 | 194 | NA | 13 | 59.25 | A |
| F2 | 196 | 61 | 14 | 61 | A |
| F66 | NA | 61 | NA | 61 | A |
| F271 | 198 | NA | 15 | 62.02 | A |
| Cm484 | NA | 62 | NA | 62 | A |
| F252 | 199 | 63 | 16 | 63 | A |
| ND283 | 204 | NA | 16 | 66.19 | B |
| NYS302 | 207 | 66 | 15 | 66 | B |
| C105 | 204 | 66 | 18 | 66 | B |
| F618 | NA | 70 | 18 | 70 | C |
| MBS847 | 210 | 71 | 18 | 71 | C |
| Lo3 | 209 | 71 | 17 | 71 | C |
| Mo17 | 217 | 73 | 17 | 73 | C |
| B73 | 217 | 74 | 21 | 74 | D |
| Lo32 | 215 | 74 | 17 | 74 | D |
| F918 | NA | 75 | 20 | 75 | D |
| HP301 | 218 | NA | 20 | 75.91 | D |
| SA24U | 220 | NA | 20 | 77.3 | D |
| F64 | 218 | 79 | 19 | 79 | D |

### Shifted sewing dates

Because we wanted all plants to be sampled at comparable developmental stage for the feeding bioassays, lines from each sewing group were sewed at different dates. At each date, six seeds per lines for all lines belonging to the same sewing groups were sewed. Below are the different sewing dates and the sewing groups that were concerned.

| Date | L | SL | SE | E |
| --- | --- | --- | --- | --- |
| 2019-10-01 | X |  |  |  |
| 2019-10-04 |  | X |  |  |
| 2019-10-07 | X |  | X |  |
| 2019-10-11 |  | X |  | X |
| 2019-10-14 | X |  | X |  |
| 2019-10-18 |  | X |  | X |
| 2019-10-21 | X |  | X |  |
| 2019-10-25 |  | X |  | X |
| 2019-10-28 |  |  | X |  |
| 2019-11-01 |  |  |  | X |

### Experimental design for feeding bioassays

Each block consisted in three batches of six maize inbred lines. Plants from the different lines were chosen to be at the same developmental stage, between five and seven visible leaf collars. Lines were randomly assigned to batches, that were launched every successive day, so that full data from one block were obtained in three days. For each batch, lines were randomly given a plate number (from *a* to *f*). A plate was filled with 50 leaf discs from the sixth leaf of the three plants from the same inbred line sewed at the same date. The developmental stage (number of visible leaf collars) and the number of days after sewing (DAS) was recorded.

The table below summarizes the experimental design.

| Line | block 1 |  |  |  |  | block 2 |  |  |  |  | block 3 |  |  |  |  |
| --- | --- | --- | --- | --- | --- | --- | --- | --- | --- | --- | --- | --- | --- | --- | --- |
|  | stage | DAS | b1.1 | b1.2 | b1.3 | stage | DAS | b2.1 | b2.2 | b2.3 | stage | DAS | b3.1 | b3.2 | b3.3 |
| F64<sup>A</sup> | V5 | 44 |  | d |  | V6 | 45 |  | c |  | V6 | 44 | d |  |  |
| SA24U<sup>A</sup> | V6 | 43 | e |  |  | V6 | 44 | a |  |  | V7 | 44 | a |  |  |
| HP301<sup>A</sup> | V6 | 43 | f |  |  | 6 | 44 | d |  |  | V6 | 44 | c |  |  |
| F918<sup>A</sup> | V5 | 43 | d |  |  | V6 | 44 | f |  |  | V7 | 44 | b |  |  |
| B73<sup>A</sup> | V6 | 43 | c |  |  | V6 | 44 | b |  |  | V7 | 44 | e |  |  |
| Lo32<sup>A</sup> | V6 | 43 | a |  |  | V6 | 44 | e |  |  | V6 | 44 | f |  |  |
| M017<sup>B</sup> | V5 | 41 |  | c |  | V5 | 42 |  | e |  | V5 | 42 |  |  | b |
| MBS847<sup>B</sup> | V5 | 41 |  | b |  | V6 | 42 |  | a |  | V6 | 41 |  | e |  |
| Lo3<sup>B</sup> | V5 | 41 |  | e |  | V5 | 42 |  | b |  | V5 | 41 |  | d |  |
| F618<sup>B</sup> | V5 | 41 |  | a |  | V6 | 42 |  | f |  | V5 | 41 |  | c |  |
| NYS302<sup>C</sup> | V5 | 38 |  | f |  | V6 | 39 |  | d |  | V6 | 41 |  | b |  |
| C105<sup>C</sup> | V6 | 37 | b |  |  | V6 | 38 | c |  |  | V6 | 39 |  | f |  |
| F252<sup>D</sup> | V5 | 35 |  |  | d | V5 | 36 |  |  | e | V5 | 39 |  | a |  |
| F271<sup>D</sup> | V5 | 35 |  |  | b | V5 | 36 |  |  | d | V5 | 36 |  |  | d |
| Cm484<sup>D</sup> | V5 | 35 |  |  | e | V5 | 36 |  |  | c | V5 | 36 |  |  | e |
| F2<sup>D</sup> | V5 | 35 |  |  | c | V5 | 36 |  |  | f | V5 | 36 |  |  | a |
| F66<sup>D</sup> | V5 | 35 |  |  | f | V5 | 36 |  |  | b | V5 | 36 |  |  | c |
| ND36<sup>D</sup> | V5 | 35 |  |  | a | V5 | 36 |  |  | a | V5 | 36 |  |  | f |

## Supplementary Data S2: Choosing the most discriminant model for CRratio

Feeding bioassays allowed to classify larvae preferences into six ordered behavioural types, named from *A* to *F*. Clearly *A* types are *consumers* that feed fast and consume all the leaf disc, while *F* types are *reluctants* that hardly feed on the leaf disc. In between, *B* to *D* behavioural profiles are intermediate. *B* and *C* mainly differ from *A* by the existence of a lag-time. *D* and *E* mainly differ from *F* by the fact that at least part of the leaf disc is consumed at the end of the experiment, but at a lower pace than in *A, B* or *C*.

In order to find significant differences between behavioural profiles, the variable CRratio was used to transform the data and analyze them on a logarithmic scale

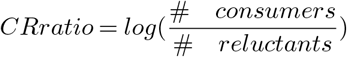

All possible grouping combination were explored and tested for the ability of the new variable CRratio to discriminate between inbred lines and replicates. For example,

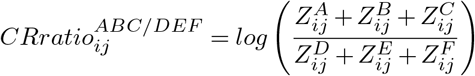

For each grouping combination, the following linear model was run:

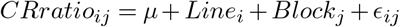

where *Line*_*i*_ is the inbred line effect, and *Block*_*j*_ is the Block effect. Summary statistics were compiled and the graph of residuals versus fitted values was checked. Results are summarized in the table below

**Table S2.** Summary statistics. pval(block) is the pvalue of the block effect. pval-line) is the pvalue of the inbred line effect. 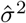 is teh residual variance. *R*2 is the adjusted model determination coefficient. *h*2 is the line heritability.

| Model | pval(block) | pval(line) | $\hat{\sigma}^2$ | R2 | h2 |
| --- | --- | --- | --- | --- | --- |
| AB/CDEF | 2e-09 | 0.0100 | 0.135 | 0.65 | 0.07 |
| A/CDEF | 3e-06 | 0.0028 | 0.278 | 0.57 | 0.19 |
| A/BCDEF | 1e-04 | 0.0014 | 0.264 | 0.54 | 0.20 |
| AB/F | 6e-04 | 0.0032 | 0.256 | 0.49 | 0.17 |
| **A/F** | **0.007** | **0.0015** | **0.528** | **0.48** | **0.40** |
| A/EF | 0.003 | 0.0074 | 0.505 | 0.43 | 0.28 |
| AB/DEF | 1e-04 | 0.0670 | 0.323 | 0.41 | 0.09 |
| A/DEF | 0.002 | 0.0301 | 0.557 | 0.38 | 0.21 |
| AB/EF | 0.005 | 0.0401 | 0.816 | 0.34 | 0.28 |
| ABC/F | 0.821 | 0.0180 | 1.222 | 0.28 | 0.54 |
| ABC/EF | 0.210 | 0.0811 | 0.869 | 0.21 | 0.22 |
| ABC/DEF | 0.061 | 0.2357 | 0.276 | 0.15 | 0.03 |

**Fig. 6.**
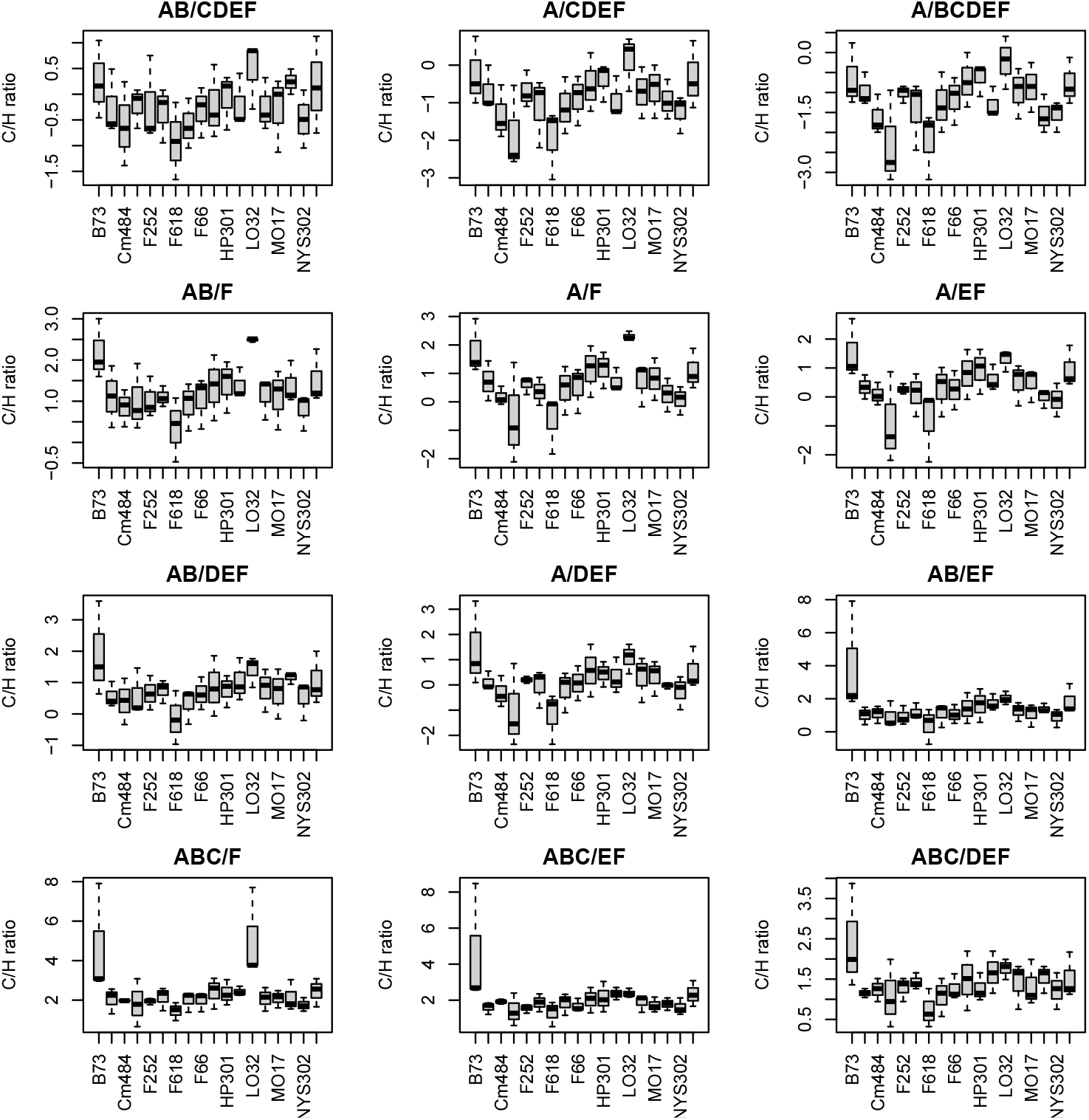
Differences between lines according to the grouping choices. Barplot representation of the variable CRratio for each maize inbred line. Grouping models have been range according to their anova R2 value.

**Fig. 7.**
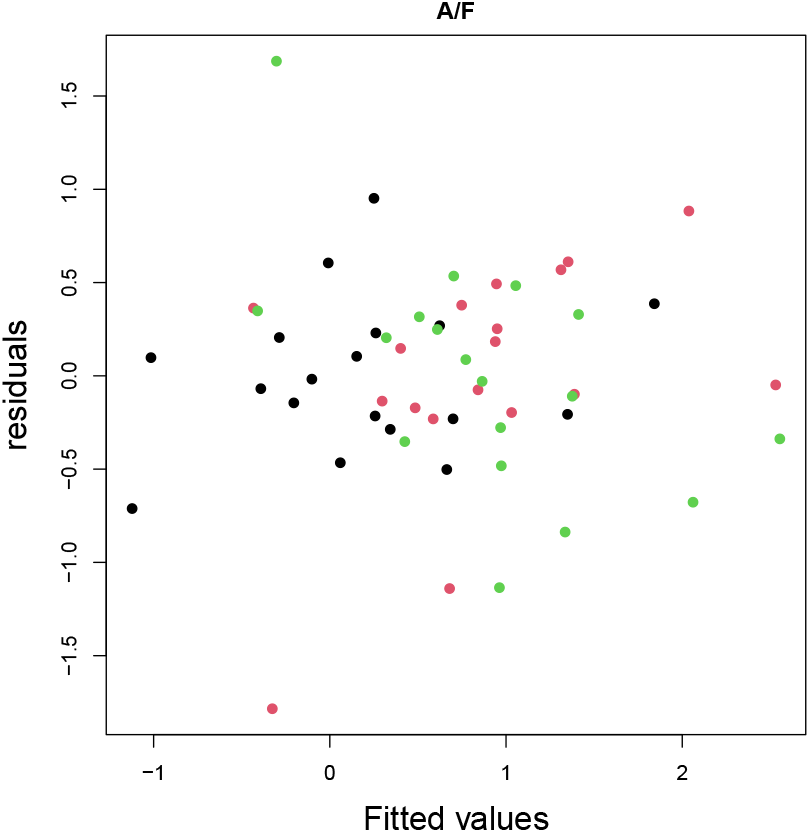
Residuals versus fitted values. The residuals plot is shown for the A/F grouping model. Colors indicate the replicates.

## Supplementary Data S3: Functional annotation of enzymes and metabolites associated with AFratio variations

We used the MetaCyc (16) and KEGG (36) databases to complete the annotation of the 15 enzymes and metabolites that were found associated to AFratio variations. Below are the names and identifiers of the molecular compounds in KEGG, PubChem and CHEBI

| Abbreviation | Name | KEGG | PubChem | ChEBI |
| --- | --- | --- | --- | --- |
| AlaAT | Alanine aminotransferase | 2.6.1.2 |  |  |
| AspAT | Aspartate aminotransferase | 2.6.1.1 |  |  |
| GS | Glutamine synthetase | 6.3.1.2 |  |  |
| NADPH-ME | malate dehydrogenase (NADP+, decarboxylating) | 1.1.1.40 |  |  |
| NADH-ME | malate dehydrogenase (oxaloacetate-decarboxylating) | 1.1.1.38 |  |  |
| NADPH-MDH | Malate dehydrogenase (NADP+) | 1.1.1.82 |  |  |
| NADH-MDH | NAD-L-malate dehydrogenase | 1.1.1.37 |  |  |
| PPDK | Pyruvate, phosphate dikinase | 2.7.9.1 |  |  |
| PFK-Pi | inorganic pyrophosphate-dependent phosphofructokinase | 2.7.1.90 |  |  |
| Glycerate | D-Glycerate | C00258 | 439194 | 32398 |
| Digalactosylglycerol | Digalactosyl-diacylglycerol | C06037 |  |  |
| Tyrosine | Tyrosine | C00082 | 6057 | 17895 |
| 5-Caffeoylquinate-trans | trans-5-O-caffeoyl-D-quinate | C00852 | 1794426 | 57644 |
| 3-Caffeoylquinate-cis | 3-Caffeoylquinate-cis |  |  |  |
| 4-Caffeoylquinate-trans | 4-Caffeoylquinate-trans |  |  |  |
| 3-Caffeoylquinate-trans | 3-Caffeoylquinate-trans |  |  |  |
| Coumaroylquinate 3061.9/345 | trans-5-O-(4-coumaroyl)-D-quinate | C12208 | 40466964 | 57575 |
| Quinate | Quinate | C00296 | 3590 | 17521 |
| Tocopherol | α-Tocopherol | C02477 | 14985 | 18145 |

Metabolic pathway databases were also used to refine the functional categories. Altogether, the 15 compounds belonged to five different main pathways that were used on Figure 4.

| Abbreviation | Class | Functional cat. | Pathway |
| --- | --- | --- | --- |
| AlaAT | Enzyme | Nitrogen assimilation | Nitrogen |
| AspAT | Enzyme | Nitrogen assimilation | Nitrogen |
| GS | Enzyme | Nitrogen assimilation | Nitrogen |
| NADPH-ME | Enzyme | Carbon fixation (mal > pyr) | Carbon |
| NADH-ME | Enzyme | Pyruvate metabolism (mal -> pyr) | Carbon |
| NADPH-MDH | Enzyme | Carbon fixation (mal -> oaa) | Carbon |
| NADH-MDH | Enzyme | Carbon fixation (mal -> oaa) | Carbon |
| PPDK | Enzyme | Carbon fixation (pyr -> pep) | Carbon |
| PFK-Pi | Enzyme | Glycolysis/Gluconeogenesis | Glycolysis/Gluconeogenesis |
| Glycerate | Carboxylic Acid | Glycerolipid metabolism | Glycerolipids |
| Digalactosylglycerol | Glycolipid | Glycerolipid metabolism | Glycerolipides |
| Tyrosine | Amino Acid | Phenylpropanoid biosynthesis | Phenylpropanoids |
| 5-Caffeoylquinate-trans | Organic Acid | Phenylpropanoid biosynthesis | Phenylpropanoids |
| 3-Caffeoylquinate-cis | Organic Acid |  | Phenylpropanoids |
| 4-Caffeoylquinate-trans | Organic Acid |  | Phenylpropanoids |
| 3-Caffeoylquinate-trans | Organic Acid |  | Phenylpropanoids |
| Coumaroylquinate 3061.9/345 | Organic Acid | Phenylpropanoid biosynthesis | Phenylpropanoids |
| Quinate | Organic acid | Phenylpropanoid biosynthesis | Phenylpropanoids |
| Tocopherol | Quinone | Quinate degradation pathway I | Phenylpropanoids |

Figure 5 shows the link between carbon fixation, nitrogen assimilation and the phenylpropanoid biosynthesis pathway. Thfigurere below shows the position of Glycerate, Digalactosilglycerol and the enzyme PFK-Pi in central carbon metabolism.

**Fig. 8.**
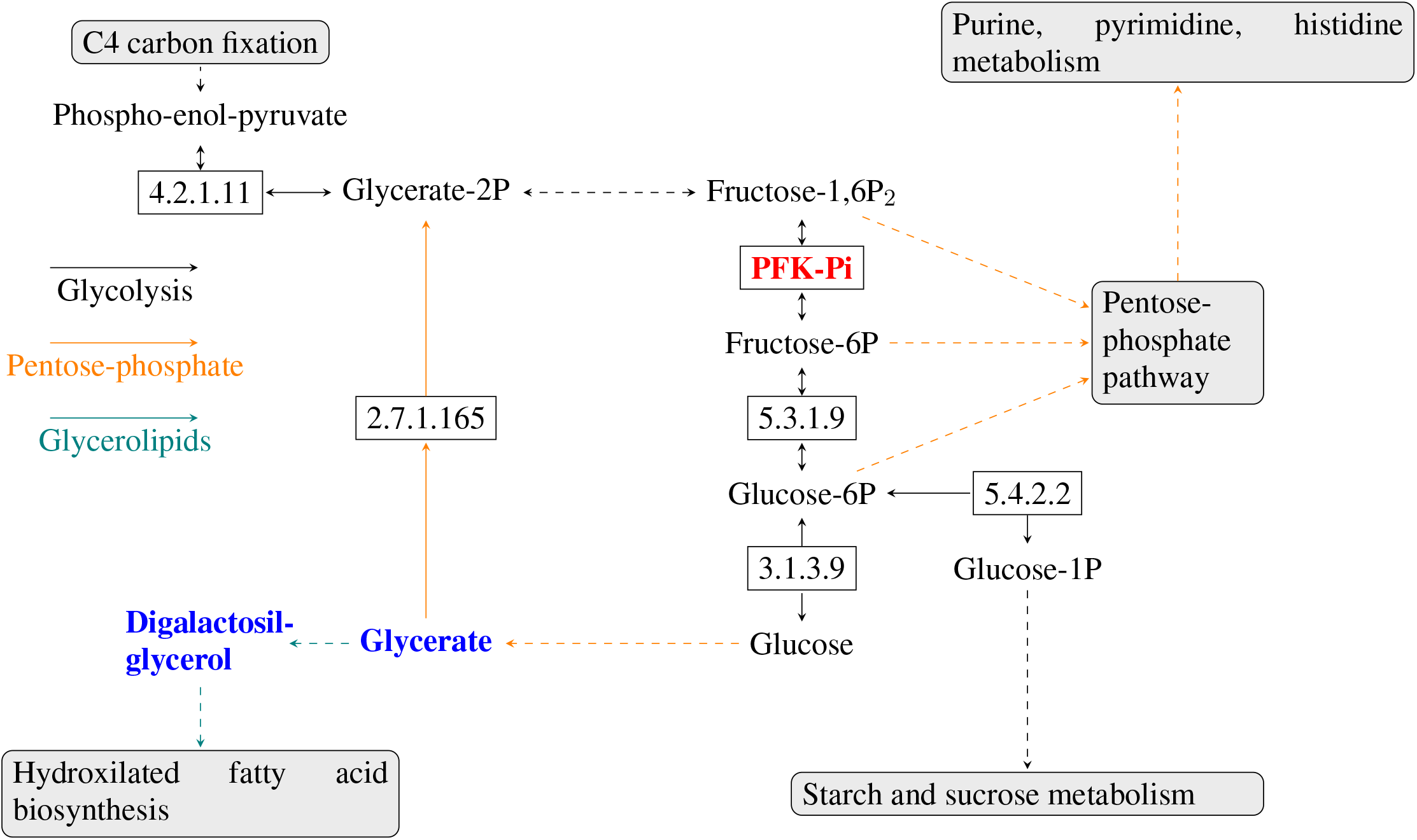
Metabolic pathways associated to maize leaf palatability: central carbon metabolism. Enzymes (rectangles) are given their EC number or their abbreviated name. Straight lines indicate a direct relation between enzymes and substrates/products and arrows the main direction of the reaction. Dashed lines indicate the link to a pathway. Line colors correspond to a pathway among glycolysis (black), pentose-phosphate (orange) or glycerolipids (green). Enzyme/metabolites colors indicate a significant positive (red) or negative (blue) association with maize leaf palatability.

